# Exposure to maternal obesity *per se* programs sex-differences in pancreatic islets of the offspring

**DOI:** 10.1101/591586

**Authors:** L M Nicholas, M Nagao, L C Kusinski, D S Fernandez-Twinn, L Eliasson, S E Ozanne

## Abstract

Maternal obesity increases type 2 diabetes (T2D) risk in the offspring. Given that nearly half of women of child-bearing age in many populations are currently overweight/obese, it is key that we improve our understanding of the impact of the *in utero*/early life environment on offspring islet function. Using a well-established mouse model of diet-induced obesity, we examined offspring islets before the onset of metabolic dysfunction. This allowed us to determine inherent changes, in males and females, which are distinct from the response of islets to an existing obesogenic, insulin resistant milieu hence identifying islet dysregulation reflecting very early manifestation of the disease before the onset of disrupted glucose homeostasis. Female offspring of obese dams displayed higher glucose-stimulated insulin secretion and mitochondrial respiration, increased expression of estrogen receptor α and decreased cleaved-caspase 3 and Bax:Bcl-2 reflecting reduced susceptibility to apoptosis. In contrast, male offspring of obese dams displayed compromised mitochondrial respiration characterised by decreased ATP synthesis-driven respiration and increased “uncoupled” respiration and reduced docked insulin granules in β-cells. Thus, maternal obesity “programs” sex-differences in offspring islet function. Islets of female but not male offspring appear primed to cope with a nutritionally-rich postnatal environment, which may reflect differences in future T2D risk.

## Introduction

More than 415 million people are living with diabetes worldwide and of this, 90% have Type 2 Diabetes (T2D) [1]. Whilst the disease has a strong genetic component, the environment also plays a significant role in its development with overweight/obesity being a major risk factor for T2D. Importantly, the rising rate of obesity also has health implications for future generations since obesity during pregnancy increases offspring obesity and T2D risk, which is thought to be mediated, at least in part, by non-genetic mechanisms (reviewed in [2]). Abnormalities in β-cell function are critical in delineating the risk of T2D because the inability of β-cells to adapt and compensate for peripheral insulin resistance leads to T2D pathogenesis [3]. In contrast, sustained β-cell adaptation is capable of preventing T2D even in the face of severe insulin resistance.

Given that in many populations, nearly half of women of child-bearing age are currently overweight/obese [4], it is important that we improve our understanding of the impact of the *in utero* and early life environment on islet function in the offspring. A number of experimental studies have examined what effect exposure to maternal obesity has on islet architecture and/or function in the offspring (reviewed in [5]). Whilst findings of reduced β-cells mass and impaired glucose-stimulated insulin secretion (GSIS) do suggest increased T2D susceptibility, it has not been possible to delineate whether these changes in islet structure/function are independent of other confounding risk factors such as increased body weight and/or adiposity in the offspring [6-10], postnatal high fat-feeding [9, 11] and the effects of ageing [8, 11].

Consequently, we have been unable to identify islet dysregulation reflecting very early manifestation of the disease before the onset of disrupted glucose homeostasis. It is, therefore, key that we identify inherent changes, which are programmed by exposure to maternal obesity *per se* and are distinct from the response of the islets to an existing obesogenic, insulin resistant milieu. This includes investigating the effects on both male and female offspring since sex affects glucose homeostasis, the pathophysiology, incidence and prevalence of T2D as well as response to therapy (reviewed in [12]). For example, using a mouse model of maternal diet-induced obesity, Samuelsson *et. al.* showed increased adiposity in both male and female offspring at six months of age, however, only male offspring developed T2D. This was characterised by higher fasting glucose, lower fasting insulin and decreased pancreatic insulin content. In contrast, these parameters were unchanged in female offspring, at the same age [13]. Thus, in the current study, using this same model we characterised pancreatic islet function in younger, metabolically healthy male and female offspring.

## Results

### Body weight and glucose tolerance are comparable between offspring of control and obese dams at eight weeks of age irrespective of sex

Offspring body weight, blood glucose and serum insulin levels (in the non-fasted state) at eight weeks of age were comparable between maternal diet groups. (Supplemental Table S1). Male offspring, however, were heavier (P<0.05) and had higher (P<0.01) serum insulin levels compared to females (Supplemental Table S1).

Glucose tolerance in both male and female offspring, as reflected by area under the curve, was also similar irrespective of maternal diet (Figures 1A and B). We found, however, that at the 15-minute time point after glucose administration, blood glucose was lower and plasma insulin levels were higher in female offspring of obese dams (Figures 1B and C).

**Figure 1:**
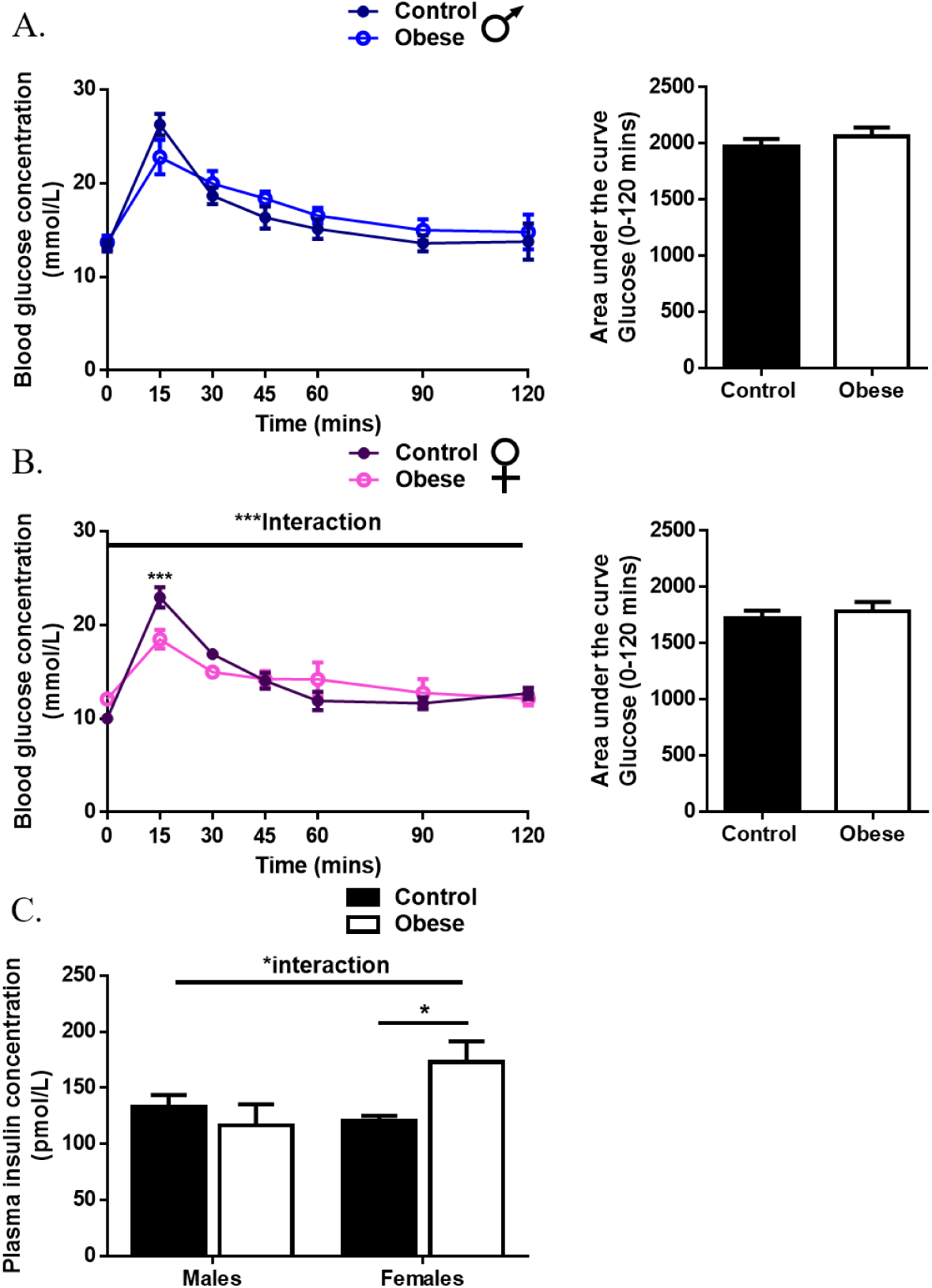
Body weight and glucose tolerance are comparable between offspring of control and obese dams at eight weeks of age irrespective of sex. Glucose excursion curve during an intraperitoneal glucose tolerance test (IPGTT) performed on seven-week-old male (A) and female (B) offspring of control and obese dams after a four hour fast. Plasma insulin concentration at the 15-minute time-point following glucose administration during an IPGTT in male and female offspring of control and obese dams (C). Data from (A) and (B) were analysed by two-way (repeated measures) ANOVA followed by Bonferonni’s multiple comparisons test. Plasma insulin concentration was analysed by two-way ANOVA followed by Tukey’s multiple comparisons test. A significant interaction between maternal diet and offspring sex indicated a sex-specific effect on the outcome measured. *P<0.05 and ***P<0.001. Males, *n*=4 – 5 mice/group and females, *n*=5 – 6 mice/group. ‘n’ represents mice from separate litters. All data are mean ± s.e.m.

### Glucose-stimulated insulin secretion is higher in female but not male offspring that were previously exposed to maternal obesity

The amount of insulin secreted by β-cells depends on the mass and function of these cells. β-cell mass was comparable between offspring of control and obese dams (Figure 2A). However, in line with having higher serum insulin levels, male offspring had increased β-cell mass compared to females. This was due to greater insulin^+^ area as well as higher absolute pancreas weight and consequently greater total pancreatic tissue area (Supplemental Figures S1A – C). In contrast, there was neither an effect of offspring sex nor exposure to maternal obesity on α-cell mass (Figure 2B).

**Figure 2:**
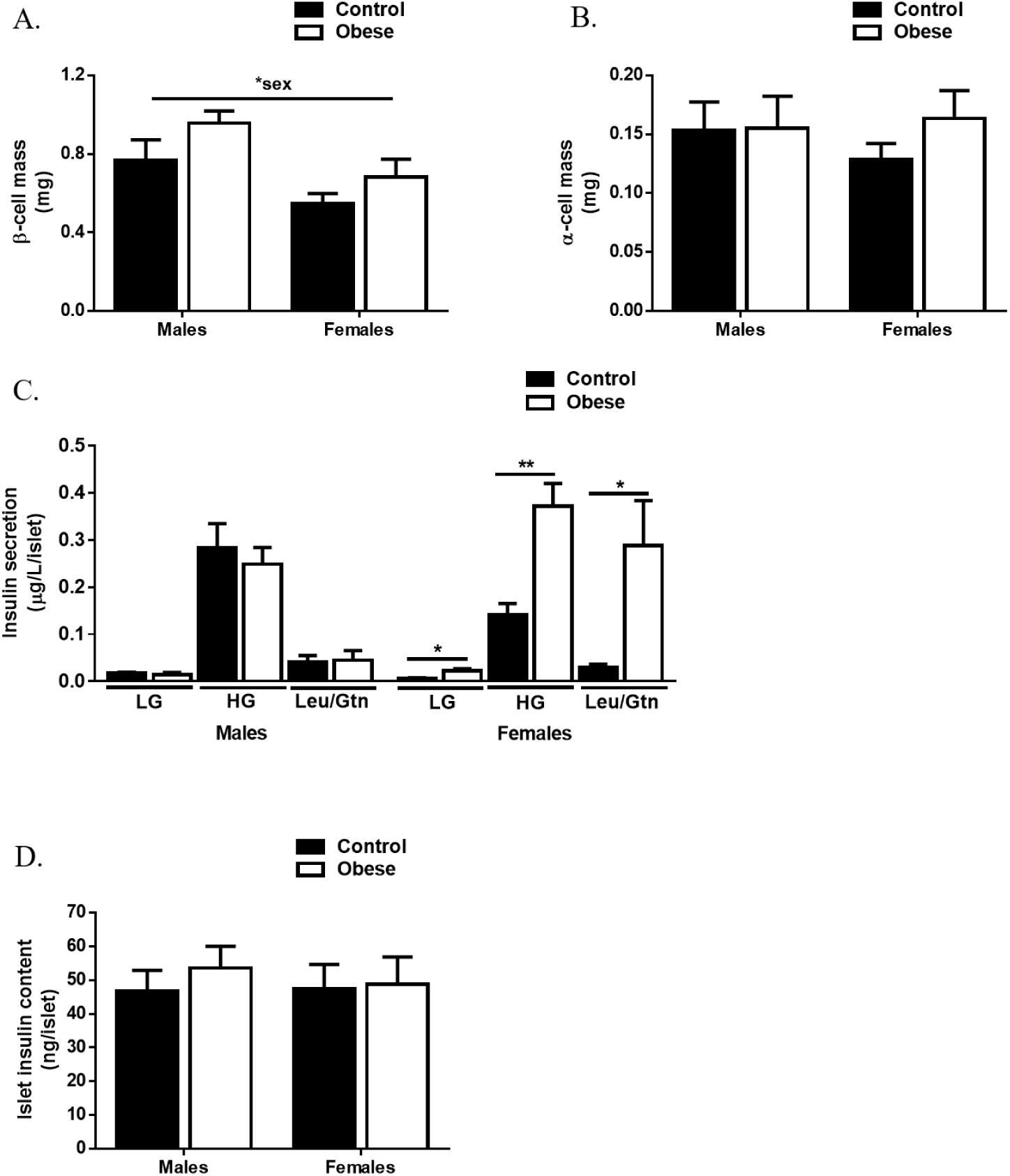
Glucose-stimulated insulin secretion is higher in female but not male offspring that were previously exposed to maternal obesity. β-cell (A) and α-cell (B) mass of offspring from control and obese dams at eight weeks of age. Low glucose (2.8mM glucose), high glucose (16.7mM glucose; HG) and 10mM leucine/glutamine stimulated insulin secretion (C). Islet insulin content (D). Experiments were performed on islets isolated from eight-week-old male and female offspring of control and obese dams. Data were analysed by two-way ANOVA followed by Tukey’s multiple comparisons test. There was a significant interaction between maternal diet and offspring sex for data relating to basal (P<0.05), glucose-stimulated (P<0.01) and leu/gtn stimulated (P<0.05) insulin secretion indicating a sex-specific effect on the outcome measured. *P<0.05 and **P<0.01. Males, *n*=6 – 9 mice/group and females, *n*=7 – 8 mice/group. ‘*n*’ represents mice from separate litters. All data are mean ± s.e.m.

Next, to determine β-cell function, we investigated GSIS in islets *ex vivo*. We found a difference between male and female offspring of obese dams. Basal, GSIS and amino acid-stimulated insulin secretion was increased in female but not male offspring of obese dams (Figure 2C). Islet insulin content was not different between any of the groups (Figure 2D).

### Sex differences in glucokinase expression and mitochondrial respiration in offspring of obese dams

To identify cellular components that may be contributing to enhanced GSIS in female offspring, we quantified protein abundance of glucose transporter-2 (GLUT2), the primary glucose transporter in rodent pancreatic islets [14]. Its expression level was not altered in either male or female offspring previously exposed to maternal obesity (Figure 3A). Subsequent phosphorylation of glucose by glucokinase (GK) is the key step controlling glycolytic flux [15]. Whilst GK abundance was also not altered in male offspring, it was reduced in female offspring of obese dams compared to controls (Figure 3B). This finding was unexpected given that the rate of glycolysis is an important determinant of GSIS from β-cells.

**Figure 3:**
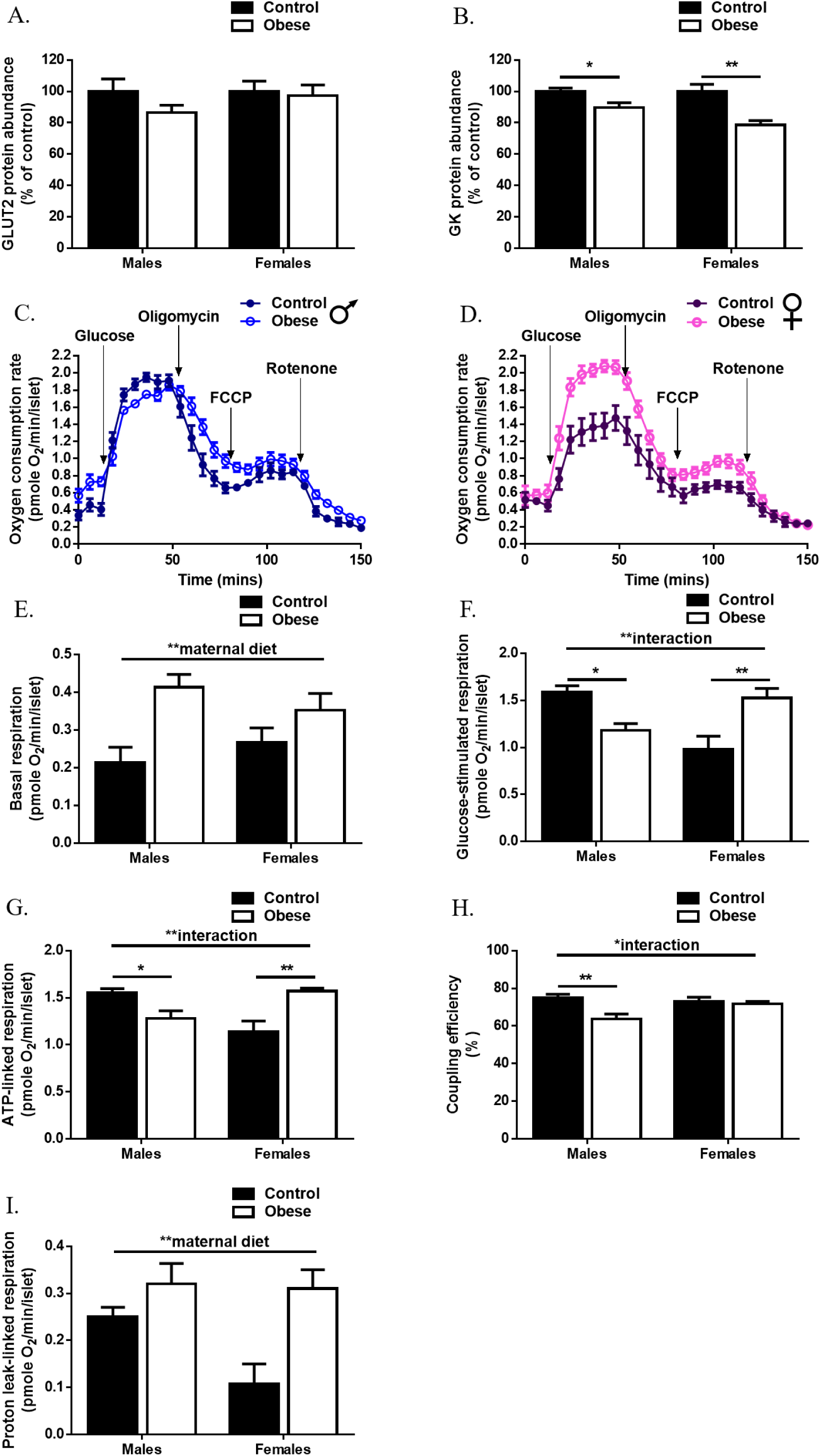
Sex differences in glucokinase expression and mitochondrial respiration in offspring of obese dams. Western blot analysis of glucose transporter 2 (A) and glucokinase (B). Changes in islet oxygen consumption rate in males (C) and females (D) offspring of control and obese dams following treatment with 16.7mM glucose, 4μg/ml oligomycin, 4μM carbonyl cyanide-4-(trifluoromethoxy) phenylhydrazone (FCCP) and 5μM rotenone. Basal (2.8mM glucose) respiration (E) glucose-simulated respiration (F) ATP-linked respiration (G) coupling efficiency (H) and proton-leak linked respiration (I). Experiments were performed on islets isolated from eight-week-old male and female offspring of control and obese dams. Data were analysed by two-way ANOVA followed by Tukey’s multiple comparisons test. A significant interaction between maternal diet and offspring sex indicated a sex-specific effect on the outcome measured. Males, *n*=4 – 7 mice/group and females, *n*=4 – 9 mice/group. ‘*n*’ represents mice from separate litters. All data are mean ± s.e.m.

As amino acid-stimulated insulin secretion was increased in female offspring of obese dams, this suggested there are pathways downstream of GK that may be activated in these islets. Distal to glucose metabolism, mitochondria metabolism couples glucose metabolism to insulin secretion in β-cells. Since leu/gtn are mitochondrial fuels, we measured cellular respiration as an indicator of mitochondrial function in islets and found sex-differences in offspring of obese dams. Whilst exposure to maternal obesity resulted in higher basal respiration irrespective of offspring sex (Figures 3C – E), glucose-stimulated respiration was different between male and female offspring of obese dams compared to controls (Figure 3F). As expected, there was a progressive rise in oxygen consumption rates upon glucose stimulation (Figures 3C and D). The magnitude of change relative to basal respiration, however, was reduced in males but increased in female offspring that were previously exposed to maternal obesity (Figure 3F).

Respiration rate is mostly composed of ATP-linked and proton leak-linked respiration [16]. The former was estimated by addition of the ATP synthase inhibitor, oligomycin. Similar to glucose-stimulated respiration, ATP-linked respiration was reduced in male offspring of obese dams (Figure 3G). Moreover, coupling efficiency, which estimates the fraction of respiration used to drive ATP synthesis was also reduced in these islets (Figure 3H). In contrast, ATP-linked respiration was increased in female offspring of obese dams (Figure 3G). Finally, we also found that proton leak-linked respiration, which would, in theory, stimulate activity of the respiratory chain was higher in islets from maternal obesity-exposed offspring (Figure 3I). This appears to be driven by a relatively large difference between female offspring of control and obese dams due to lower proton leak-linked respiration in the control group.

### Only female offspring of obese dams have increased expression of mitochondrial and nuclear encoded components of the electron transport chain

Mitochondrial respiration/function is critically dependent on mitochondrial DNA, the genes that it encodes (most notable of which are those that are part of the respiratory chain complexes) and the nuclear-encoded constituents of these complexes. mRNA expression of some mitochondrial-encoded components of the respiratory complexes was significantly increased in maternal obesity-exposed female offspring (Figure 4A). This increase was not due to higher levels of the mitochondrial transcription factor, TFAM (Figures 4B) but may be partly due to increased mitochondria as evidenced by higher mitochondrial area density (Figure 4C) and mitochondrial DNA content, which tended to be increased (P<0.06; Figure 4D). In addition to mitochondrial genes, mRNA expression of *Sdha* (Complex II) (Figure 4A) and protein abundance of SDHB (Complex II), UQCRC2 (Complex III) and ATP5A (Complex V), which are nuclear-encoded was also increased (Figure 4E).

**Figure 4:**
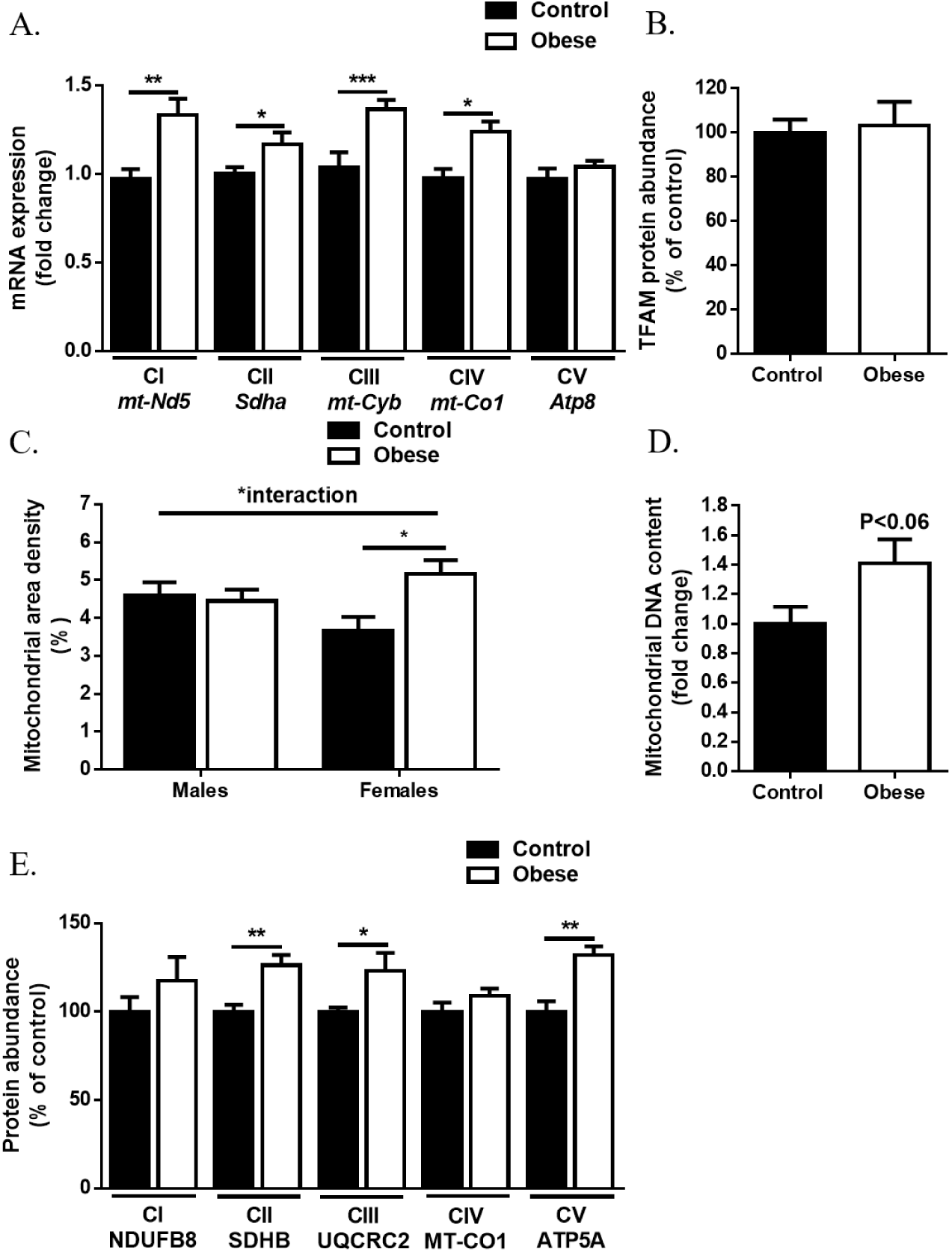
Only female offspring of obese dams have increased expression of mitochondrial and nuclear encoded components of the electron transport chain. qRT-PCR analysis of mRNA expression of mitochondrial (*mt-Nd5, mt-Cyb, mt-Co1, mt-Atp8*) and nuclear (*Sdha*) encoded components of the electron transport chain (A). Western blot analysis of mitochondrial transcription factor A (TFAM) (B). Mitochondrial area density (C) Mitochondrial DNA content (D). Western blot analysis of mitochondrial (MT-CO1) and nuclear (NFUFB8, SDHB, UQCRC2 and ATP5A) encoded components of the electron transport chain (E). Experiments were performed on islets isolated from eight-week-old female offspring of control and obese dams. Data were analysed independently by unpaired Student’s t-test (Control versus Obese). *P<0.05, **P<0.01 and ***P<0.001. Females, *n*=5 – 10 mice/group. ‘*n*’ represents mice from separate litters. All data are mean ± s.e.m.

Taken together, our results suggest that the higher mitochondrial respiration in female offspring of obese dams could be due to higher levels of respiratory complex proteins. The converse is not the case for male offspring since these parameters were unchanged in male offspring of obese dams compared to controls (Supplemental Figures S2A – D and 4C).

### Increased ROS in islets from both male and female offspring of obese dams but increased expression of antioxidant enzyme only in female islets

Mitochondria are a major source of intracellular reactive oxygen species (ROS) production [17]. We, therefore, measured ROS levels in islets under basal and glucose-stimulated conditions to determine whether changes in mitochondrial respiration had an impact on their levels. Basal ROS levels tended to be slightly higher (P<0.06; Figure 5A) in islets of maternal obesity-exposed offspring, which could be the result of increased basal respiration. ROS levels remained higher in these offspring in response to glucose stimulation, this time reaching statistical significance (Figure 5B). Thus, in the case of male offspring of obese dams, ROS levels are elevated even with reduced mitochondrial respiration.

**Figure 5:**
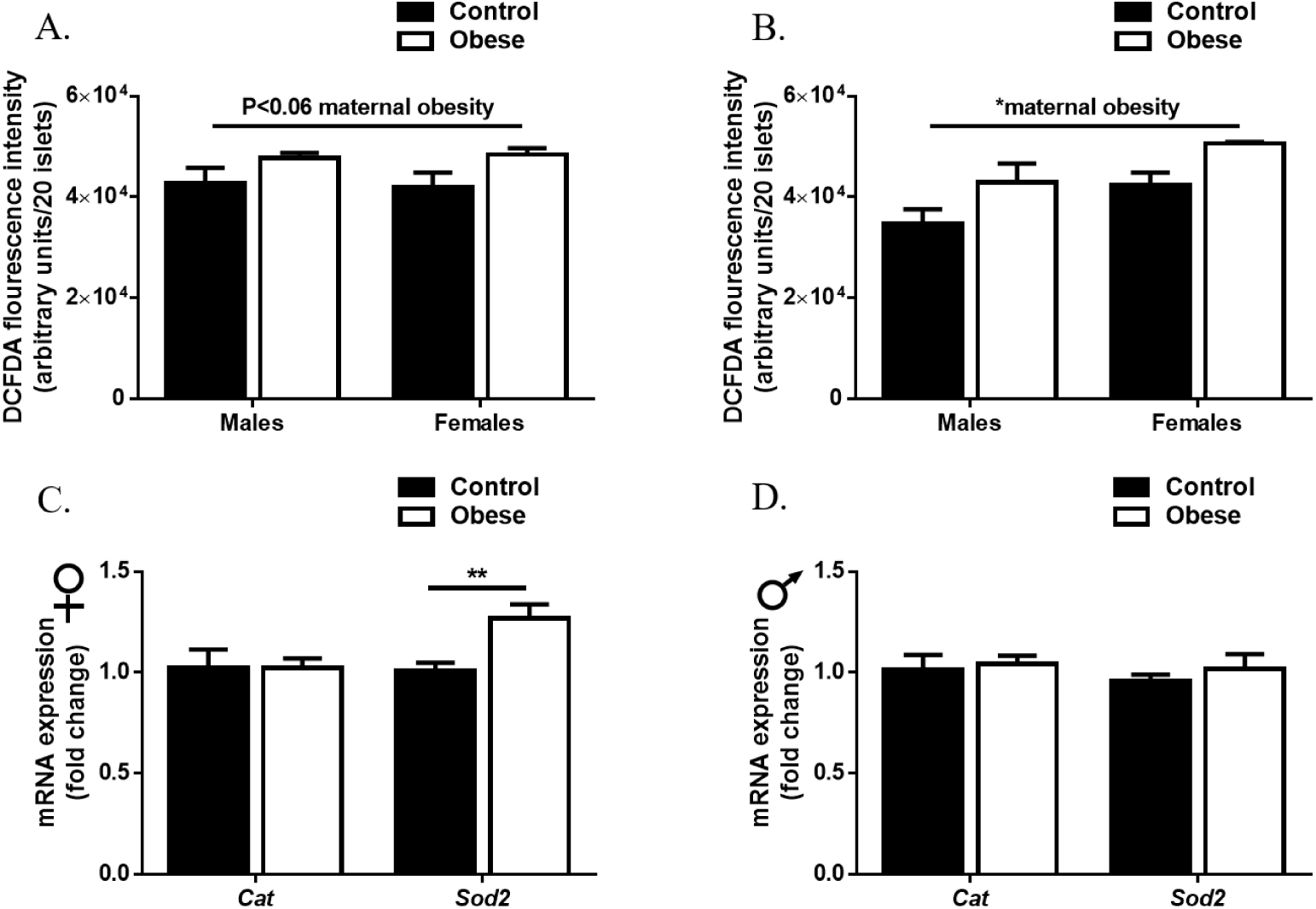
Increased ROS in islets from both male and female offspring of obese dams but increased expression of antioxidant enzyme only in female islets. ROS levels measured by DCFDA fluorescence intensity, following incubation with 2.8mM glucose (A) and 16.7mM glucose (B). qRT-PCR analysis of *Cat* and *Sod2* mRNA expression in female (C) and male (D) offspring. Experiments were performed on islets isolated from eight-week-old female offspring of control and obese dams. Data relating to DCFDA fluorescence intensity were analysed by two-way ANOVA followed by Tukey’s multiple comparisons test. Male and female offspring data relating to mRNA expression were analysed independently by unpaired Student’s t-test (Control versus Obese). *P<0.05 and **P<0.01. Males, *n*=5 – 8 mice/group and females, *n*=4 – 9 mice/group. ‘*n*’ represents mice from separate litters. All data are mean ± s.e.m.

β-cells are particularly vulnerable to oxidative stress due to the high levels of oxygen consumption required for insulin secretion coupled with the relatively low levels of antioxidant enzymes [18]. Female but not male offspring of obese dams displayed increased *Sod2* (but not *Cat)* mRNA expression (Figures 5C and D) suggesting that islets from these male offspring may be more vulnerable to oxidative stress.

### Reduction in docked insulin granules in β-cells from male offspring that were previously exposed to maternal obesity

One of the pre-requisites of insulin exocytosis is docking of secretory vesicles at the plasma membrane. Thus, we examined β-cell morphology by electron microscopy and quantified the density of granules including those that were docked. The density of insulin granules was reduced in β-cells of offspring that were exposed to maternal obesity (Figure 6A). When we examined docked granules, however, we found that the surface density was significantly decreased in islets from male but not female offspring of obese dams (Figure 6B). Furthermore, the number of granules in the first fraction beneath the plasma membrane (0-0.19µm) was reduced whilst those in the second fraction (0.19 – 0.38µm from the plasma membrane) were increased in these mice (Figure 6C). These results suggest that granules may be stacking in the second fraction (which is considered to be the reserve pool) due to dysfunctional docking in β-cells of these mice (Representative micrographs: Supplemental Figure S3).

**Figure 6:**
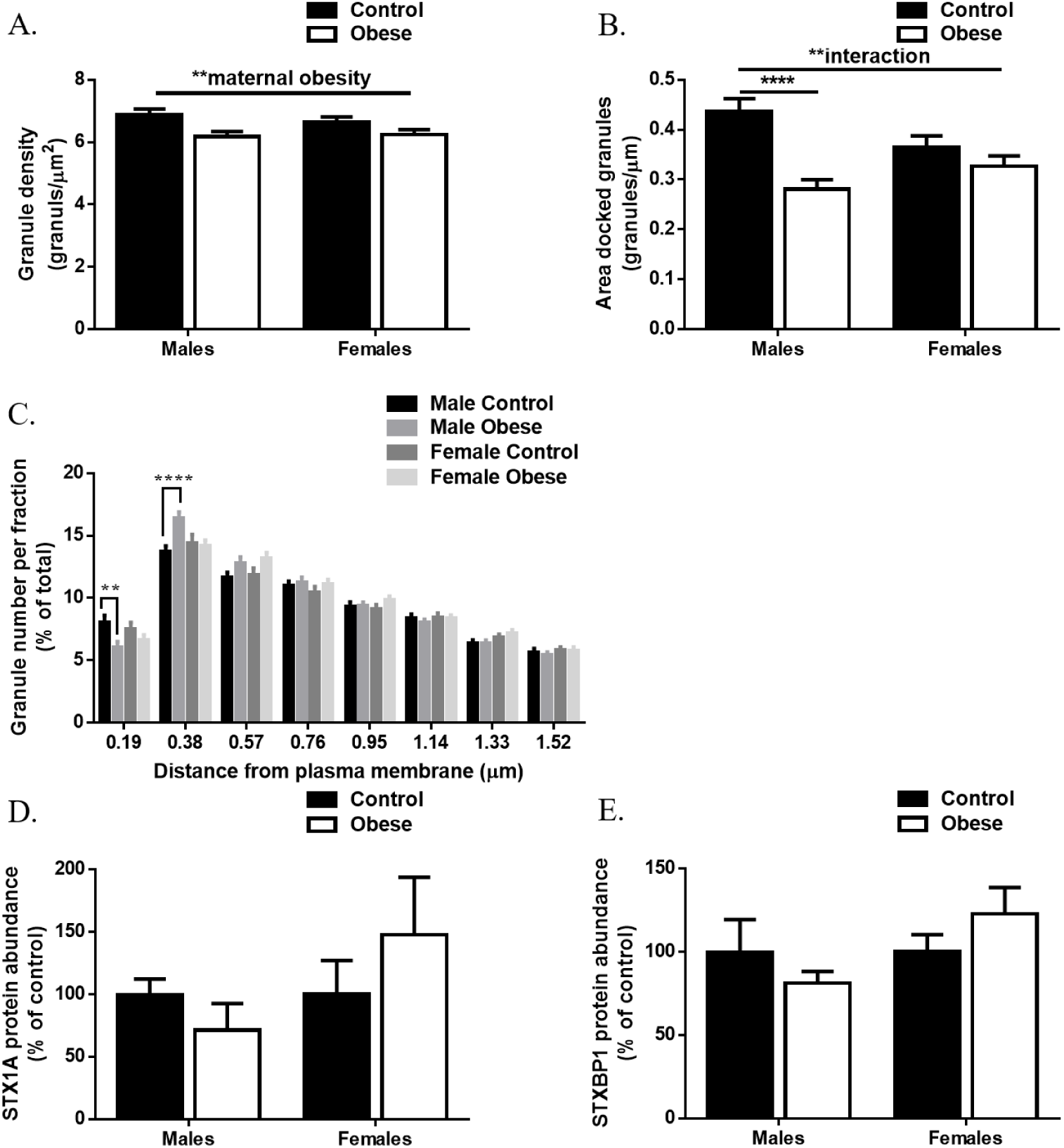
Reduction in docked insulin granules in β-cells from male offspring that were previously exposed to maternal obesity. Transmission electron microscopy analysis of insulin granule density (A) and docked granules estimated by the surface density (B). Relative distribution of granules at distance fractions from the plasma membrane (C). Granules were defined as docked if their distance from the plasma membrane was 0.19µm i.e. half the size of the mean granule diameter. Western blot analysis of Syntaxin 1A (D) and syntaxin binding protein 1 (E). Experiments were performed on islets isolated from eight-week-old female offspring of control and obese dams. Data relating to electron microscopy images were analysed by two-way ANOVA followed by Tukey’s multiple comparisons test. Analyses based on individual animals (*n*=2 – 3 mice/group) showed no statistically significant difference within the groups. Thus, cell-based analyses (*n*= 40 – 59 cells/group) was performed. Male and female offspring data relating to protein expression were analysed independently by unpaired Student’s t-test (Control versus Obese). **P<0.01 and ****P<0.0001. Males, *n*=4 – 5 mice/group and females, *n*=5 – 6 mice/group. ‘*n*’ represents mice from separate litters. All data are mean ± s.e.m.

Docking requires the plasma membrane SNARE protein syntaxin-1A (STX1A) and its binding partner, munc-18 (STXBP1) [19]. The reduction in docked granules observed in male offspring of obese dams, however, was not due to altered abundance of these proteins (Figures 6D and E). STX1A and STXBP1 protein abundance was also comparable between female offspring of control and obese dams (Figures 6D and E).

### Islets from female offspring of obese dams may be protected from dysfunction by increased estrogen receptor α and reduced susceptibility to apoptosis

Females are protected from β-cell death and hyperglycemia in most rodent models of T2D [12, 20]. This has been attributed, in part, to the actions of 17β-estradiol via estrogen receptor α (ERα) and its role in preventing apoptosis [21-23]. In our model, ERα protein abundance was higher in female offspring of obese dams (Figure 7A). Furthermore, expression of cleaved (activated) caspase-3, the key mediator of the apoptotic cascade in mammalian cells [24] was decreased in female offspring of obese dams compared to controls (Figure 7B). In contrast, cleaved caspase-3 abundance and *Casp3* mRNA expression was increased in male offspring of obese versus control dams (Figures 7B and C).

**Figure 7:**
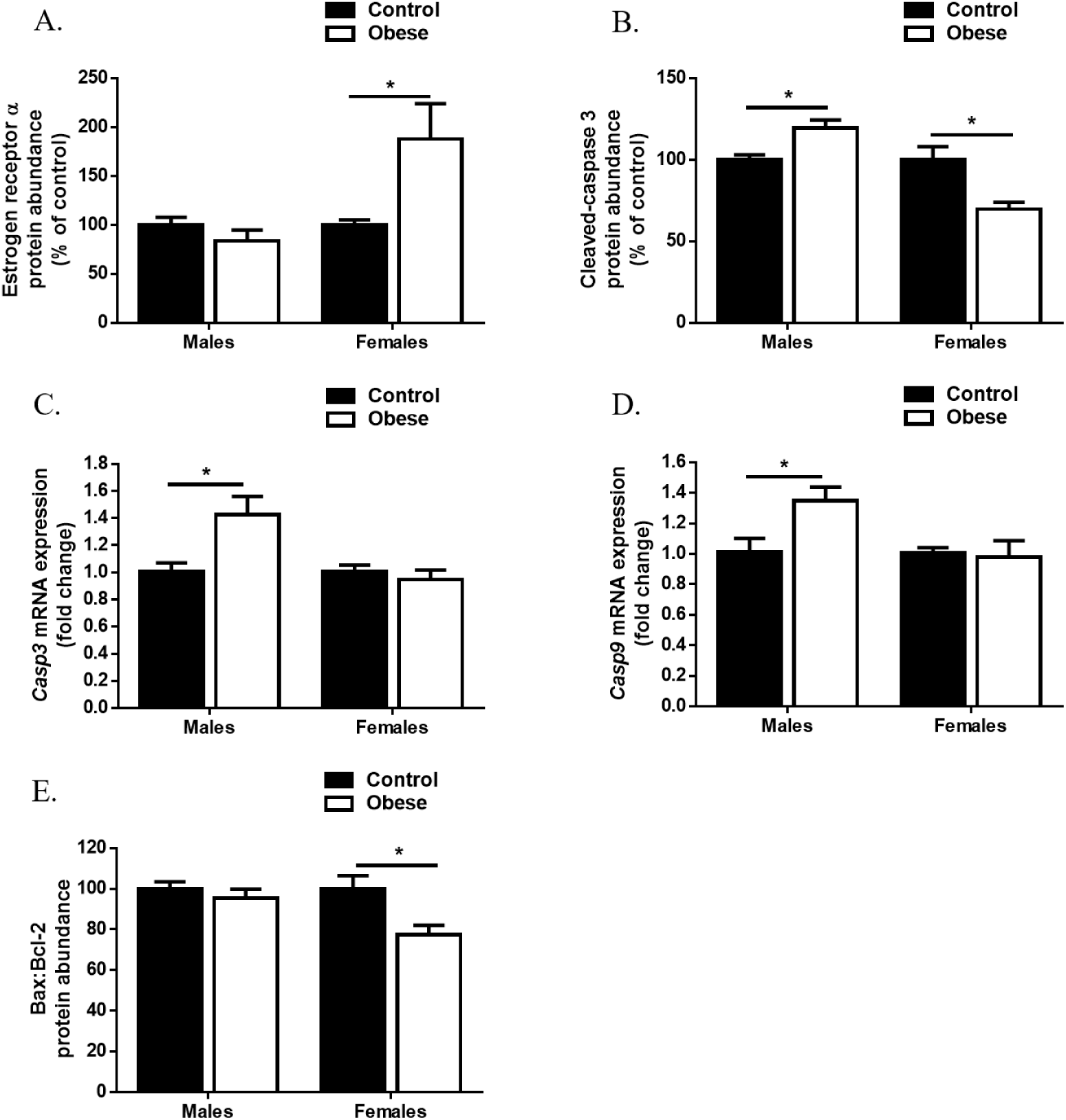
Islets from female offspring of obese dams may be protected from dysfunction by increased estrogen receptor α and reduced susceptibility to apoptosis. Western blot analysis of estrogen receptor α (A) and cleaved-caspase 3 (B) protein abundance. qRT-PCR analysis of *Casp3* and *Casp9* mRNA expression in male (C) and female (D) offspring. Western blot analysis of Bax/Bcl-2 ratio. Experiments were performed on islets isolated from eight-week-old female offspring of control and obese dams. Males and females were analysed independently by unpaired Student’s t-test (Control versus Obese). *P<0.05 and **P<0.01. Males, *n*=4 – 6 mice/group and females, *n*=4 – 9 mice/group. ‘*n*’ represents mice from separate litters. All data are mean ± s.e.m.

Considering the mitochondrial phenotype observed in this study, we sought to determine whether the mitochondrial intrinsic apoptosis pathway could be contributing to changes in caspase-3 activity. mRNA expression of *Casp9*, the initiator caspase involved in this pathway was unchanged in females but significantly higher in male offspring of obese dams (Figure 7D). The regulation of this pathway also occurs through members of the Bcl-2 family of proteins, which govern mitochondrial membrane permeability and can be either pro-or anti-apoptotic [25]. The ratio of pro-apoptotic Bax to anti-apoptotic Bcl-2 has been described as a cellular ‘rheostat’ of apoptosis sensitivity and can profoundly influence the ability of a cell to respond to an apoptotic signal [26]; a lower Bax:Bcl-2 reflects reduced susceptibility to an apoptotic stimuli. Whist we found no differences in protein levels of Bax and Bcl-2 between offspring of maternal diet groups (Supplemental Figures S4A and B), Bax:Bcl-2 was lower in female offspring of obese dams compared to controls (Figure 7E) suggesting a down-regulation of the mitochondrial apoptosis pathway in these offspring.

## Discussion

Whilst there are known genetic risk factors that may be transferred from mother to child, which explain future risk of obesity and T2D in her offspring, the picture is far from complete. Studies in humans that have controlled for shared genetics by comparing siblings discordant for a particular exposure e.g. gestational diabetes mellitus [27, 28] or siblings born before versus after maternal bariatric surgery [29] found that exposed individuals had a greater risk for diabetes and obesity than unexposed siblings. Given that in humans a mother’s genes and environment co-exist, rodent models of maternal obesity across pregnancy and lactation using obesogenic diets that resemble the human situation have been key in our understanding of the non-genetic transfer of metabolic disease risk from mother to offspring.

A number of studies have suggested that increased T2D susceptibility in developmentally programmed offspring is due to altered pancreatic islet architecture and/or function (reviewed in [5]). This is the first study, however, to outline the sex-specific changes in islet function in offspring born to obese dams that are present before and are therefore independent of offspring obesity, insulin resistance and ageing. This is important as it allows us to elucidate islet processes that are most vulnerable to dysfunction thus leading to higher T2D risk in these offspring as a consequence of exposure to maternal obesity.

It has been shown previously using the same model of maternal diet-induced obesity, that whilst both male and female offspring develop obesity and insulin resistance with age, female offspring of obese dams are less susceptible to T2D compared to males at six months of age [13]. Findings from the current study suggest that islets from obese female offspring are primed to handle a nutritionally-rich postnatal environment by up-regulating mitochondrial respiration leading to higher GSIS. This is reflected by lower blood glucose and higher serum insulin levels 30 minutes after glucose administration during and IPGTT. Therefore, although these islets are being “overworked” from a young age, they appear to have protective mechanisms in place to cope with the demands of increasing adiposity and insulin resistance. For example, reduced GK expression could act to temper glycolysis in the face of increased mitochondrial metabolism so as to maintain a healthy redox balance [30]. Increased proton-leak linked respiration may also not necessarily be damaging (especially since respiration coupling efficiency is maintained). Mitochondrial superoxide production is steeply dependent on the protonmotive force across the inner mitochondrial membrane. Thus, increased proton leak may act to minimize oxidative damage by moderating the protonmotive force and, therefore, ROS production [31]. Furthermore, increased ERα levels could also have a positive influence on these islets by preserving mitochondrial function [32] and protecting them from hyperglycemia associated β-cell apoptosis [21-23].

Females appear to be protected from β-cell death in most rodent models of T2D [12, 20]. In the case of exposure to maternal diet-induced obesity/high fat-feeding and future T2D risk in the offspring, Yokomizo and colleagues showed that islets from female offspring were better to able to compensate and adapt to a high fat diet (HFD) in postnatal life compared to male offspring [9]. Interestingly, in a study of maternal obesity using the agouti viable yellow (A^vy^) mouse, (aged) female offspring developed glucose intolerance and had reduced GSIS following postnatal HFD compared to males [11]. This is in contrast to a study by Li and colleagues who found that the latent predisposition to metabolic disease in offspring of A^vy^ dams was more prominent in males who were significantly heavier and developed glucose intolerance and insulin resistance after only three weeks of HFD [33]. It should be noted that in contrast to most models of maternal *diet-induced* obesity, (A^vy^) mice are normoglycemic during pregnancy.

In this study, islets from male offspring that were exposed to maternal obesity appear to not fare as well as female offspring. This manifests as sub-optimal mitochondrial respiration; there is decreased respiration to drive ATP synthesis, the efficiency of this process is compromised and there is increased “uncoupled” respiration. The latter, however, may also act to alleviate ROS production in these islets. Unlike in females, these changes could not be attributed to changes in mitochondrial DNA content or the expression of genes encoding components of the electron transport chain. The mitochondria in islets of male offspring seem particularly vulnerable to insults *in utero*/early life; male offspring that experienced intrauterine growth restriction followed by postnatal catch-up growth also showed impaired mitochondrial function and increased ROS production prior to the onset of diabetes [34]. This suggests there may be common responses operating as a result of suboptimal nutrition *in utero*. There also appears to be a problem with docking of insulin granules in male offspring of obese dams. This could be due to reduced ATP levels [35] owing to reduced ATP-linked respiration in these islets. Whilst not yet detrimental to GSIS at this age, these changes could mean that their islets are more vulnerable to metabolic insult/stress and may partly explain why male offspring that were exposed to maternal obesity are more at risk of developing T2D.

In summary, the current findings suggest that known sex-differences in T2D susceptibility in offspring as a consequence of exposure to maternal obesity could, at least in part, be driven by differences in islet function. Females, in response to nutritional cues from the mother signalling a nutrient-rich environment, prime islet development to thrive in this environment postnatally. In contrast, males may respond to these cues in a manner that minimises the risk of neonatal hypoglycaemia and maximises neonatal survival. However, this becomes maladaptive in later life following the onset of obesity and insulin resistance in these male offspring.

## Experimental procedures

### Animal model

This research was regulated under the UK Home Office Animals (Scientific Procedures) Act 1986 following ethical review by the University of Cambridge Animal Welfare and Ethical Review Board. The model has been described in detail previously [36]. Briefly, four-week-old female C57BL/6 mice were fed *ad libitum* either a standard control chow (7% simple sugars, 3% fat [w/w]) RM1 diet or a highly palatable energy-rich obesogenic diet (10% simple sugars, 20% animal fat [w/w]) and sweetened condensed milk (55% simple sugars, 8% fat, 8% protein [w/w] (Nestle) fortified with mineral and vitamin mix (AIN93G), for six weeks before mating for first pregnancy. The first litter was culled post-weaning. This first pregnancy ensured the mice were proven breeders. Mice were then re-mated for a second pregnancy. Dams were maintained on their respective diets throughout both pregnancies and lactation periods. Litter size was standardised to six pups on postnatal day two. Offspring from ‘Control’ and ‘Obese’ groups were weaned onto RM1 and remained on this diet until the end of the study. Intraperitoneal glucose tolerance test (IPGTT) was performed on seven-week-old offspring and they were sacrificed at eight weeks of age. Prior to sacrifice, tail blood glucose was measured using a blood glucose analyser (AlphaTRAK). Blood was taken by cardiac puncture for serum insulin analysis. In all relevant experiments, insulin concentration was determined by ELISA (Mercodia).

### Intraperitoneal glucose tolerance test (IPGTT)

Following a four hour fast, a bolus of glucose (1g/kg) was injected into the intraperitoneal cavity. Blood glucose measurements were made using a blood glucose analyser (AlphaTRAK). Tail blood was also collected at 0, 15 and 30 mins into glass micro-haematocrit capillary tubes with sodium heparin (Hirschmann-Laborgeräte) for plasma insulin analysis. Area under the curve for glucose was calculated using the trapezoidal rule.

### Pancreatic islet isolation

Pancreas was perfused with Collagenase P solution (Roche) through the common bile duct, harvested and digested at 37°C for 18 min. Islets were hand-picked under a stereo microscope and incubated at 37°C overnight in a humidified atmosphere of air and 5% CO_2_ in RPMI-1640 medium supplemented with 10% fetal bovine serum, 100U/ml penicillin and 100μg/ml streptomycin sulphate.

### Islet insulin secretion

Islets were incubated in Krebs-Ringer bicarbonate buffer (KRBB) [37] at pH 7.2 with 2mg/ml bovine serum albumin (BSA) and 2.8mM glucose for 1 hr at 37°C. Next, groups of 20 islets were incubated at 37°C for 1 h in KRBB containing 2.8mM glucose, 16.7mM glucose and 10mM leu/gtn.

### Islet insulin content

Islets were washed in PBS, mixed with ethanol/hydrochloric acid and sonicated at 4°C. After centrifugation, the supernatant was stored at – 20°C until assayed.

### Oxygen consumption assay

Oxygen consumption rates were measured by the XF24 Extracellular Flux Analyzer (Agilent). Groups of 50 islets (in triplicate) were pre-incubated in KRBB with 2mg/ml BSA and 2.8mM glucose for 30 mins at 37°C. Respiration was measured in the presence of 16.7 mM glucose, oligomycin, FCCP and rotenone as previously described [38]. Calculations of basal, glucose-stimulated, ATP-linked and proton leak-linked respiration and coupling efficiency was carried out according to [39].

### Quantification of mRNA expression using quantitative real-time-PCR (qRT-PCR)

Total RNA was extracted from islets (Qiagen) and reverse transcribed into cDNA (Fermentas). qRT-PCR was performed using the SYBR Green system (Thermo Fisher Scientific). mRNA expression was determined by the ΔΔCt method and normalized to the expression of *Rplp0* and *Hprt* (for females) and *Actb* and *Hprt* (for males). Primer sequences are available in Supplemental Table 2.

### Quantification of protein abundance using Western blotting

Islets (*n*≥4 mice/sex/treatment group) were lysed in radioimmunoprecipitation assay buffer with protease inhibitor cocktail (Sigma, St. Louis, MO, USA). Lysates were subjected to SDS-PAGE and blotted using antibodies as detailed in Supplemental Table 3. Male and female samples were run on separate gels. Membranes were sectioned and some sections were stripped (Restore™ Western Blot Stripping Buffer, Thermo Fischer Scientific) and re-probed to maximise the amount of data obtained from each Western blot. Image Lab software version 5.2.1 (Bio-Rad, Hercules, CA, USA) was used to quantify the density of specific bands. Images of all Western blots are included in Supplemental Figure S4.

### Mitochondrial DNA content

Genomic and mitochondrial DNA was extracted using DNeasy Blood and Tissue Kit (Qiagen). DNA was quantified using Quant-iT™ PicoGreen™ dsDNA Reagent (Thermo Fisher Scientific) and equal quantities amplified by qRT-PCR using primers for nuclear (Rplp0) and mitochondrial DNA (mt-Nd5). Mitochondrial DNA content, relative to nuclear DNA was determined by the following equations: ΔC_T_ = (nuclear DNA C_T_ – mitochondrial DNA C_T_) and relative mitochondrial DNA content = 2 × 2ΔC_T_ [40].

### Transmission electron microscopy

Groups of 40-50 isolated islets were fixed in Millonig’s buffer and post-fixed in 1.0% osmium tetroxide, dehydrated and embedded in AGAR 100 (Oxfors Instruments Nordiska AB). 70-90nm ultrathin sections were cut, mounted and contrasted before being examined in JEM 1230 electron microscope (JEOL-USA, Inc.) Electron micrographs of at least 40 different β-cells from 2-3 mice were taken for each group. Granules (large dense-core vesicles) were defined as docked when the centre of the granule was located within 190 nm (a half-length of the mean granule diameter) from the plasma membrane. The distance between the center of the granule and the plasma membrane was calculated using an in-house software programmed in MatLab 7 (MathWorks, Natick, MA, USA) [41]. Granule density was estimated according to the procedures described by DeHoff and Rhines [42] and were normalized to granule diameter assuming spherical geometry [41].

### Tissue processing and immunofluorescence

Paraffin embedded pancreas was exhaustively sectioned at 5μm thickness, 200μm apart. Sections were incubated with primary antibody against insulin (Dako) and glucagon (Bioss Antibodies Inc). The appropriate fluorescent-dye conjugated secondary antibodies were used for identifying β- and α-cells (Jackson ImmunoResearch). Digital images were obtained using a Zeiss Axioscan Z1 Slide Scanner and HALO™ image analysis platform (Indica Labs) was used to calculate immune-positive and total tissue area. β-cell and α-cell mass was calculated using the following formula: β/α-cell mass (mg) = insulin^+^/glucagon^+^ area (μm^2^)/total pancreatic tissue area (μm^2^) × pancreas weight (mg).

## Statistical analyses

All calculations were performed in IBM SPSS Statistics 23. ‘*n*’ represents the number of litters. All data are presented as mean ± SEM of the indicated number of litters. Data relating to gene expression, protein abundance, mitochondrial DNA content, and AUC_glucose_ were analysed using unpaired Student’s t-test. Blood glucose and insulin levels during IPGTT was measured by two-way (repeated measures) ANOVA followed by Bonferonni’s multiple comparisons test (*Note:* each sex was analysed separately). All other data were analysed by two-way ANOVA followed by Tukey’s multiple comparisons test. A significant interaction between maternal diet and offspring sex indicated a sex-specific effect on the outcome measured. A probability level of 5% (P<0.05) was taken to be significant.

## Acknowledgements

We would like to thank Thomas J Ashmore and Claire Custance for expert technical assistance.

## Author contributions

L.M.N. conceived and performed experiments, analysed data and wrote the manuscript. M.N. performed experiments and analysed data. L.C.K and D.S.F.T performed experiments. L.E. provided expertise and feedback. L.M.N., L.E. and S.E.O. interpreted the data. L.M.N and S.E.O are the guarantors of this work and, as such, had full access to all the data in the study and take responsibility for the integrity of the data and the accuracy of the data analysis. All authors read, commented on and approved the final version of the manuscript.

## References

1. Zheng, Y., S.H. Ley, and F.B. Hu, Global aetiology and epidemiology of type 2 diabetes mellitus and its complications. Nature Reviews Endocrinology, 2017. 14: p. 88.

2. Nicholas, L.M., et al., The early origins of obesity and insulin resistance: timing, programming and mechanisms. Int J Obes (Lond), 2016. 40(2): p. 229–38.

3. Kahn, S.E., R.L. Hull, and K.M. Utzschneider, Mechanisms linking obesity to insulin resistance and type 2 diabetes. Nature, 2006. 444(7121): p. 840–846.

4. Branum, A.M., S.E. Kirmeyer, and E.C. Gregory, Prepregnancy Body Mass Index by Maternal Characteristics and State: Data From the Birth Certificate, 2014. Natl Vital Stat Rep, 2016. 65(6): p. 1–11.

5. Elsakr, J.M. and M. Gannon, Developmental programming of the pancreatic islet by in utero overnutrition. Trends Dev Biol, 2017. 10: p.79–95.

6. Graus-Nunes, F., et al., Pregestational maternal obesity impairs endocrine pancreas in male F1 and F2 progeny. Nutrition, 2015. 31(2): p. 380–7.

7. Srinivasan, M., et al., Maternal high-fat diet consumption results in fetal malprogramming predisposing to the onset of metabolic syndrome-like phenotype in adulthood. Am J Physiol Endocrinol Metab, 2006. 291(4): p. E792–9.

8. Taylor, P.D., et al., Impaired glucose homeostasis and mitochondrial abnormalities in offspring of rats fed a fat-rich diet in pregnancy. American Journal of Physiology - Regulatory Integrative and Comparative Physiology, 2005. 288(1 57-1).

9. Yokomizo, H., et al., Maternal high-fat diet induces insulin resistance and deterioration of pancreatic beta-cell function in adult offspring with sex differences in mice. Am J Physiol Endocrinol Metab, 2014. 306(10): p. E1163–75.

10. Zambrano, E., et al., Decreased basal insulin secretion from pancreatic islets of pups in a rat model of maternal obesity. J Endocrinol, 2016. 231(1): p. 49–57.

11. Han, J., et al., Long-term effect of maternal obesity on pancreatic beta cells of offspring: reduced beta cell adaptation to high glucose and high-fat diet challenges in adult female mouse offspring. Diabetologia, 2005. 48(9): p. 1810–8.

12. Gannon, M., et al., Sex differences underlying pancreatic islet biology and its dysfunction. Mol Metab, 2018. 15: p. 82–91.

13. Samuelsson, A.M., et al., Diet-induced obesity in female mice leads to offspring hyperphagia, adiposity, hypertension, and insulin resistance: A novel murine model of developmental programming. Hypertension, 2008. 51(2): p. 383–392.

14. Thorens, B., et al., Cloning and functional expression in bacteria of a novel glucose transporter present in liver, intestine, kidney, and beta-pancreatic islet cells. Cell, 1988. 55(2): p. 281–90.

15. Meglasson, M.D. and F.M. Matschinsky, New perspectives on pancreatic islet glucokinase. Am J Physiol, 1984. 246(1 Pt 1): p. E1–13.

16. Divakaruni, A.S., et al., Methods in enzymology. Methods in enzymology, 2014. 547C): p. 309.

17. Andreyev, A.Y., Y.E. Kushnareva, and A.A. Starkov, Mitochondrial metabolism of reactive oxygen species. Biochemistry (Mosc), 2005. 70(2): p. 200–14.

18. Lei, X.G. and M.Z. Vatamaniuk, Two tales of antioxidant enzymes on beta cells and diabetes. Antioxid Redox Signal, 2011. 14(3): p. 489–503.

19. Gandasi, N.R. and S. Barg, Contact-induced clustering of syntaxin and munc18 docks secretory granules at the exocytosis site. Nat Commun, 2014. 5: p. 3914.

20. Louet, J.F., C. LeMay, and F. Mauvais-Jarvis, Antidiabetic actions of estrogen: insight from human and genetic mouse models. Curr Atheroscler Rep, 2004. 6(3): p. 180–5.

21. Alonso-Magdalena, P., et al., Pancreatic insulin content regulation by the estrogen receptor ER alpha. PLoS One, 2008. 3(4): p. e2069.

22. Kilic, G., et al., The islet estrogen receptor-alpha is induced by hyperglycemia and protects against oxidative stress-induced insulin-deficient diabetes. PLoS One, 2014. 9(2): p. e87941.

23. Le May, C., et al., Estrogens protect pancreatic beta-cells from apoptosis and prevent insulin-deficient diabetes mellitus in mice. Proc Natl Acad Sci U S A, 2006. 103(24): p. 9232–7.

24. Hui, H., et al., Role of caspases in the regulation of apoptotic pancreatic islet beta-cells death. J Cell Physiol, 2004. 200(2): p. 177–200.

25. Cory, S. and J.M. Adams, The Bcl2 family: regulators of the cellular life-or-death switch. Nat Rev Cancer, 2002. 2(9): p. 647–56.

26. Oltvai, Z.N., C.L. Milliman, and S.J. Korsmeyer, Bcl-2 heterodimerizes in vivo with a conserved homolog, Bax, that accelerates programmed cell death. Cell, 1993. 74(4): p. 609–19.

27. Dabelea, D., et al., Intrauterine exposure to diabetes conveys risks for type 2 diabetes and obesity: a study of discordant sibships. Diabetes, 2000. 49(12): p. 2208–11.

28. Lawlor, D.A., P. Lichtenstein, and N. Langstrom, Association of maternal diabetes mellitus in pregnancy with offspring adiposity into early adulthood: sibling study in a prospective cohort of 280,866 men from 248,293 families. Circulation, 2011. 123(3): p. 258–65.

29. Smith, J., et al., Effects of maternal surgical weight loss in mothers on intergenerational transmission of obesity. J Clin Endocrinol Metab, 2009. 94(11): p. 4275–83.

30. Wu, L., et al., Oxidative stress is a mediator of glucose toxicity in insulin-secreting pancreatic islet cell lines. J Biol Chem, 2004. 279(13): p. 12126–34.

31. Divakaruni, A.S. and M.D. Brand, The regulation and physiology of mitochondrial proton leak. Physiology (Bethesda), 2011. 26(3): p. 192–205.

32. Zhou, Z., et al., Estrogen receptor alpha protects pancreatic beta-cells from apoptosis by preserving mitochondrial function and suppressing endoplasmic reticulum stress. J Biol Chem, 2018. 293(13): p. 4735–4751.

33. Li, C.C., et al., Maternal obesity and diabetes induces latent metabolic defects and widespread epigenetic changes in isogenic mice. Epigenetics, 2013. 8(6): p. 602–11.

34. Simmons, R.A., I. Suponitsky-Kroyter, and M.A. Selak, Progressive accumulation of mitochondrial DNA mutations and decline in mitochondrial function lead to beta-cell failure. J Biol Chem, 2005. 280(31): p. 28785–91.

35. Eliasson, L., et al., Rapid ATP-dependent priming of secretory granules precedes Ca(2+)-induced exocytosis in mouse pancreatic B-cells. J Physiol, 1997. 503 (Pt 2): p. 399–412.

36. Martin-Gronert, M.S., et al., Altered hepatic insulin signalling in male offspring of obese mice. J Dev Orig Health Dis, 2010. 1(3): p. 184–91.

37. Nicholas, L.M., et al., Mitochondrial transcription factor B2 is essential for mitochondrial and cellular function in pancreatic β-cells. Molecular metabolism, 2017. 6(7): p. 651–663.

38. Malmgren, S., et al., Tight coupling between glucose and mitochondrial metabolism in clonal beta-cells is required for robust insulin secretion. J Biol Chem, 2009. 284(47): p. 32395–404.

39. Brand, M.D. and D.G. Nicholls, Assessing mitochondrial dysfunction in cells. Biochem J, 2011. 435(2): p. 297–312.

40. Rooney, J.P., et al., PCR based determination of mitochondrial DNA copy number in multiple species. Methods Mol Biol, 2015. 1241: p. 23–38.

41. Olofsson, C.S., et al., Fast insulin secretion reflects exocytosis of docked granules in mouse pancreatic B-cells. Pflugers Arch, 2002. 444(1-2): p. 43–51.

42. Dehoff, R.T. and F.N. Rhines, Determination of the number of particles per unit volume from measurements made on random plane sections: the general cylinder and the ellipsoid. Transactions of the Metallurgical Society of AIME, 1961. 221: p. 975–982.

