## Supplemental Information for "Exposure to maternal obesity *per se* programs sex-differences in pancreatic islets of the offspring"

**Supplemental Table 1: Body weight and serum glucose and insulin concentrations in male and female offspring of control and obese dams.** Blood was collected from fed animals at eight weeks of age. Data were analysed by two-way ANOVA followed by Tukey's multiple comparisons test. *P<0.05 and **P<0.01 for an effect of offspring sex. Males, *n*=4 – 6 mice/group and females, *n*=4 – 9 mice/group. '*n*' represents mice from separate litters. All data are mean ± s.e.m.

| | **Control** | | **Obese** | |
|---|---|---|---|---|
| | **Male** | **Female** | **Male** | **Female** |
| **Body weight (g)** | 25.3 ± 0.7* | 20.6 ± 0.4 | 25.3 ± 0.9* | 20.5 ± 0.5 |
| **Blood glucose concentration (mmol/L)** | 13.2 ± 1.2 | 11.5 ± 0.8 | 12.7 ± 1.1 | 10.9 ± 0.5 |
| **Serum insulin concentration (µg/L)** | 245 ± 31** | 124 ± 17 | 257 ± 29** | 166 ± 34 |

**Supplemental Table 2: Oligonucleotide primer sequences for qRT-PCR analysis**

| **Gene Name (GenBank accession no.)** | **Forward** | **Reverse** |
|---|---|---|
| *Actb* (NM_007393.5) | 5’ GCCCTGAGGCTCTTTTCCAG3’ | 5’ TGCCACAGGATTCCATACCC3’ |
| *Hprt* (NM_013556.2) | 5’ GAGAGCGTTGGGCTTACCTC3’ | 5’ ATCGCTAATCACGACGCTGG3’ |
| *Rplp0* (NM_007475.5**)** | 5’GAAAAGTTCTTGATCCCCAATGC3’ | 5’ TGTGACTGGTCCACAATTCCTT3’ |
| *mt-Nd5* (NC_005089.1) | 5’ACGAAAATGACCCAGACCTC3’ | 5’ AGATGACAAATCCTGCAAAGATG3’ |
| *mt-Cytb* (NC_005089.1) | 5’CCCACCCCATATTAAACCCG3’ | 5’GAGGTATGAAGGAAAGGTATAAGGG3’ |
| *mt-Co1* (NC_005089.1) | 5’TCCCAGATATAGCATTCCCACG3’ | 5’ACTGTTCATCCTGTTCCTGC3’ |
| *mt-Atp8* (NC_005089.1) | 5’GCCACAACTAGATACATCAACATG3’ | 5’TGGTTGTTAGTGATTTTGGTGAAG3’ |
| *Sdha* (NM_023281.1) | 5’TTACAAAGTGCGGGTCGATGA3’ | 5’TGTTCCCCAAACGGCTTCTT3’ |
| *Cat* (NM_009804.2) | 5’AGCGACCAGATGAAGCAGTG3’ | 5’TCCGCTCTCTGTCAAAGTGTG3’ |

| | | |
|---|---|---|
| *Sod2* (NM_013671.3) | 5’CAGACCTGCCTTACGACTATGG3’ | 5’CTCGGTGGCGTTGAGATTGTT3’ |
| *Casp3* (NM_001284409.1) | 5’TGGTGATGAAGGGGTCATTTATG3’ | 5’TTCGGCTTTCCAGTCAGACTC3’ |
| *Casp9* (NM_001277932.1) | 5’TCCTGGTACATCGAGACCTTG3’ | 5’AAGTCCCTTTCGCAGAAACAG3’ |

**Supplemental Table 3: Antibodies used for Western blotting**

| **Protein** | **Company** | **Catalogue No.** |
|---|---|---|
| GLUT2 | Abcam | ab54460 |
| NDUFB8 | Abcam | ab110413 |
| SDHB | Abcam | ab110413 |
| UQCRC2 | Abcam | ab110413 |
| MT-CO1 | Abcam | ab110413 |
| ATP5A | Abcam | ab110413 |
| TFAM | Abcam | ab47517 |
| β-Tubulin | Abcam | ab15568 |
| BAX | Cell Signalling Technology | #2772 |
| BCL-2 | Cell Signalling Technology | #3498 |
| Cleaved caspase 3 | Cell Signalling Technology | #9662 |
| STX1A | Synaptic Systems | 110 111 |
| STXBP1 | Synaptic Systems | 116 002 |
| ERα | Santa Cruz Biotechnology | sc-8002 |
| GCK | Santa Cruz Biotechnology | sc-7908 |

**Figure S1: Male offspring had higher insulin⁺ area, absolute pancreas weight and total pancreatic tissue area compared to females, irrespective of maternal diet.** Insulin⁺ area (A), absolute pancreas weight (B) and total pancreatic tissue area (C) in eight-week-old male and female offspring of control and obese dams. Data were analysed by two-way ANOVA followed by Tukey's multiple comparisons test. *P<0.05. Males, *n*=4 mice/group and females, *n*=6 mice/group. '*n*' represents mice from separate litters. All data are mean ± s.e.m.

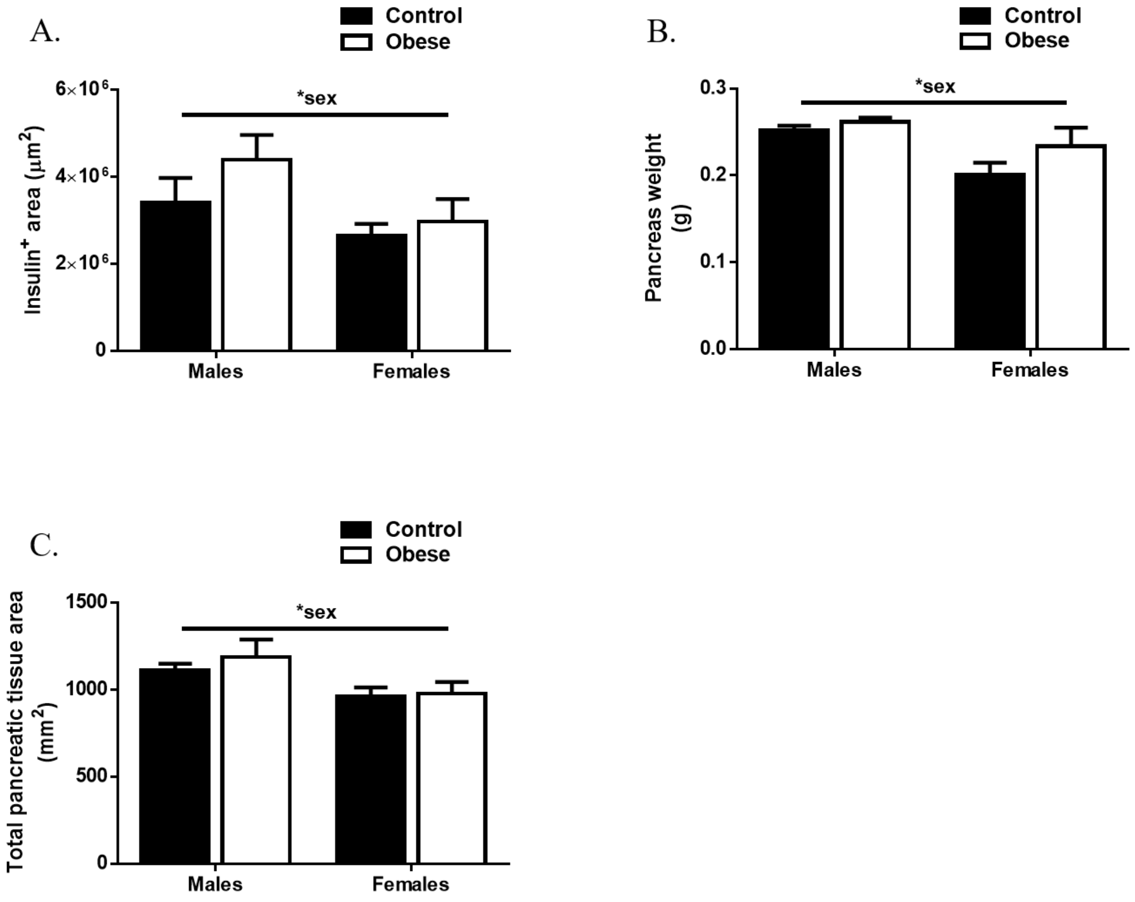

**Figure S2: Exposure of male offspring to maternal obesity had minimal impact on mitochondrial DNA content and expression of mitochondrial and nuclear-encoded components of the electron transport chain.** Mitochondrial DNA content (A). qRT-PCR analysis of mRNA expression of mitochondrial (*mt-Nd5*, *mt-Cyb*, *mt-Co1*, *mt-Atp8*) and nuclear (*Sdha*) encoded components of the electron transport chain (B). Western blot analysis of mitochondrial transcription factor A (TFAM) (C) and mitochondrial (MT-CO1) and nuclear (NFUFB8, SDHB, UQCRC2 and ATP5A) encoded components of the electron transport chain (D). Experiments were performed on islets isolated from eight-week-old male mice. Data were analysed independently by unpaired Student's t-test (Control versus Obese). *P<0.05, **P<0.01 and ***P<0.001. *n*=5 – 8 mice/group. '*n*' represents mice from separate litters. All data are mean ± s.e.m.

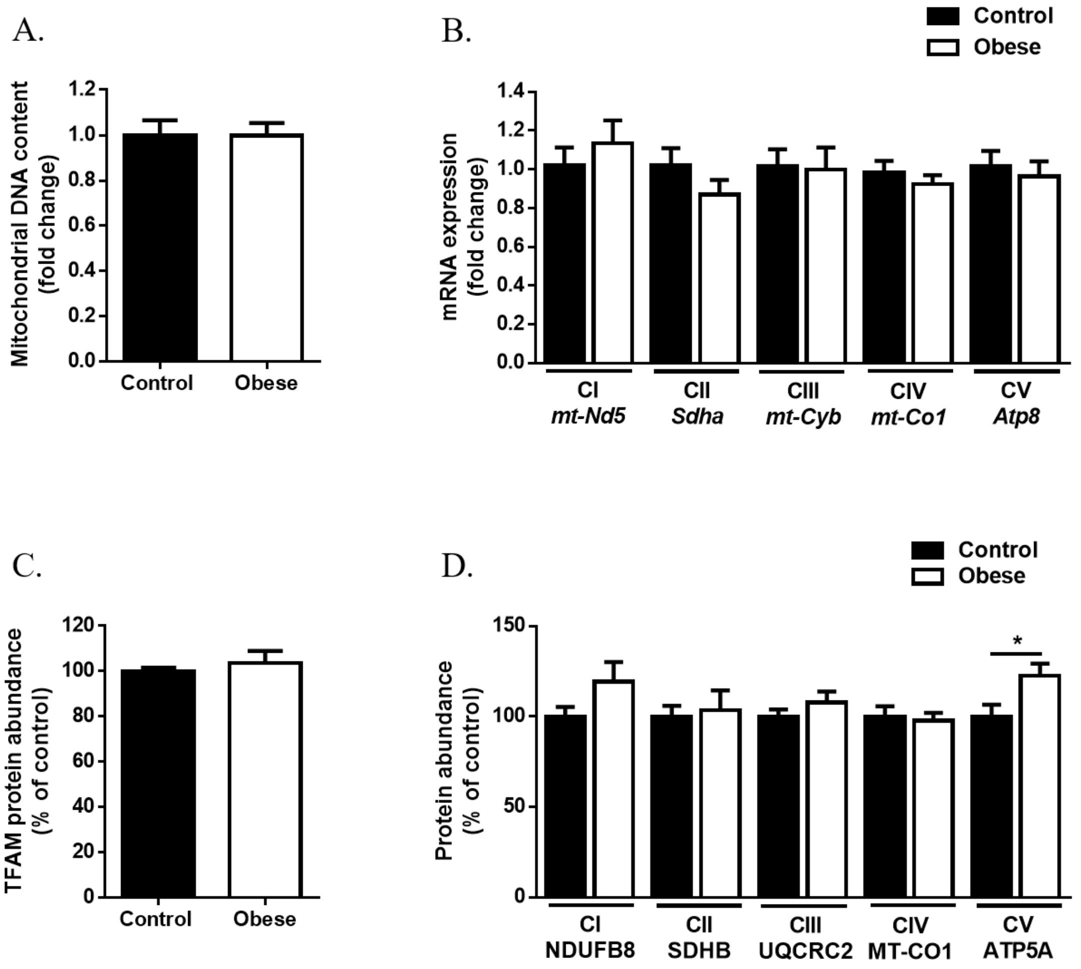

**Figure S3: Representative transmission electron microscopy micrographs.** β-cell from male offspring of control dam (A) male offspring of obese dam (B) female offspring of control dam (C) and female offspring of obese dam (D).

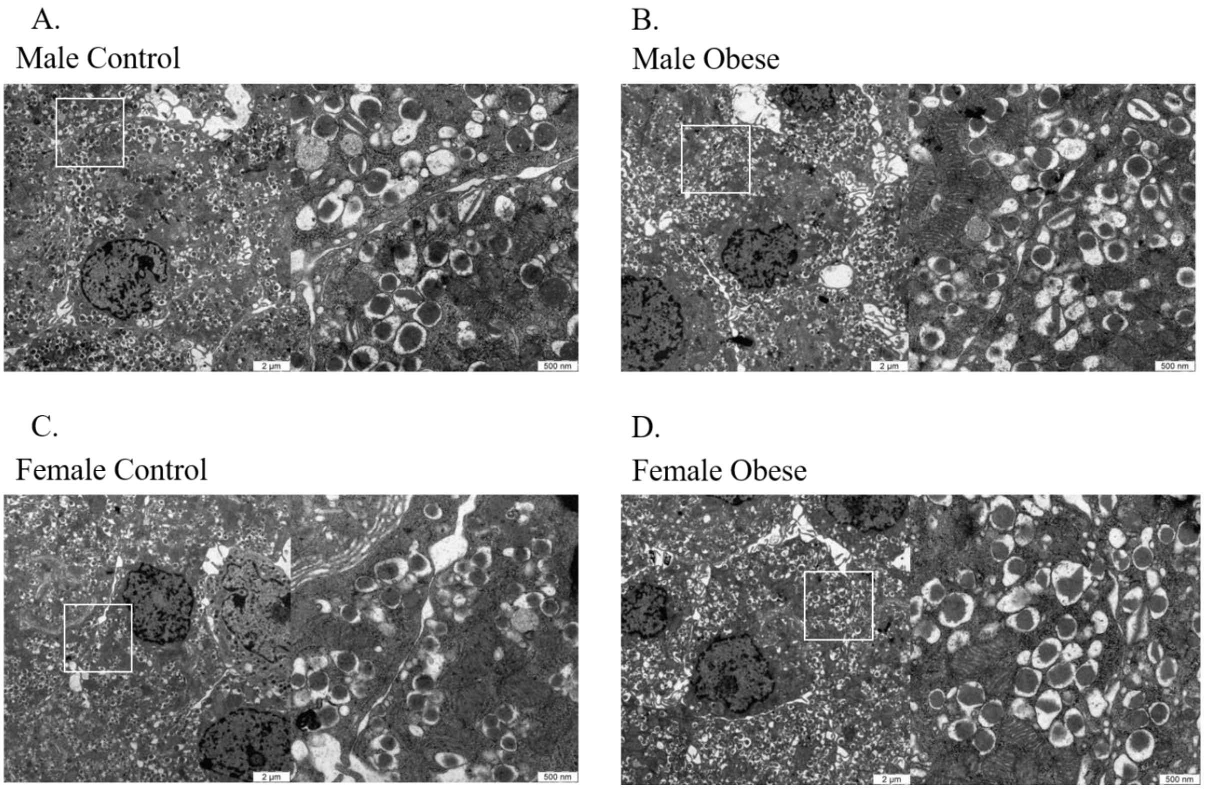

**Figure S4: Exposure to maternal obesity had no impact on the expression of pro-apoptotic Bax or anti-apoptotic Bcl-2.** Western blot analysis and Bax (A) and Bcl-2 (B). Experiments were performed on islets isolated from eight-week-old offspring. Male and female offspring were analysed independently by unpaired Student's t-test (Control versus Obese). Males, *n*=4 – 5 mice/group and females, *n*=4 – 5 mice/group. '*n*' represents mice from separate litters. All data are mean ± s.e.m.

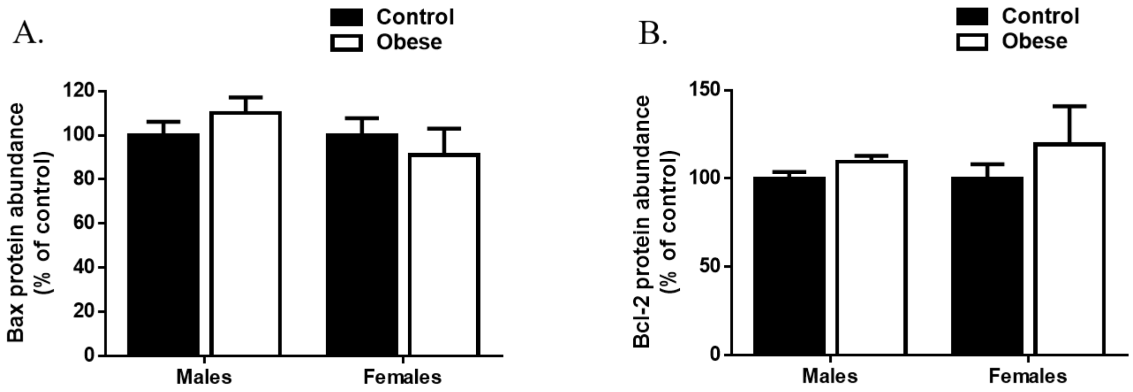

**Figure S5: Images of Western blots used for quantification of protein abundance**

**Glucose transporter 2 (Females)**

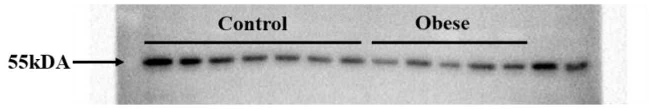


**β-tubulin (Females)**

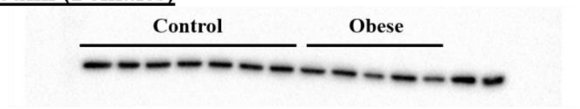


**Glucose transporter 2 (Males)**

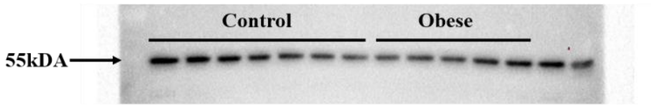


**β-tubulin (Males)**

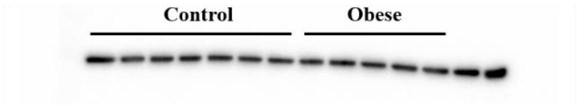


**Glucokinase (Females)**

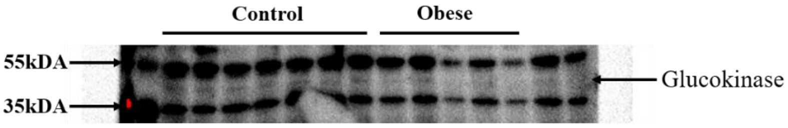


**β-tubulin (Females)**

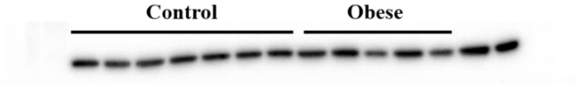


**Glucokinase (Males)**

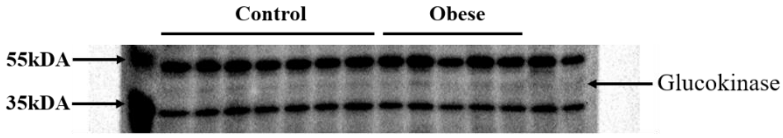


**β-tubulin (Males)**

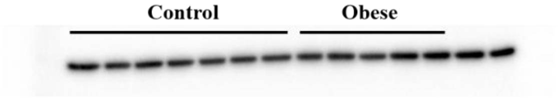

**Mitochondrial transcription factor A (Mitochondrial) (Females)**

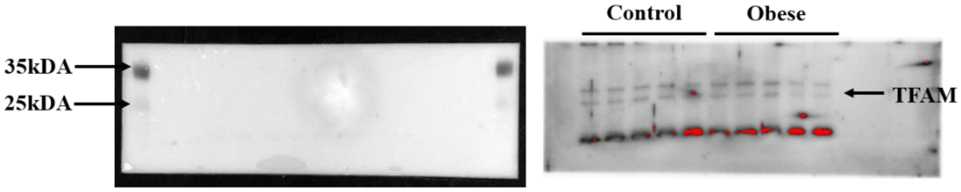


**β-tubulin (Females)**

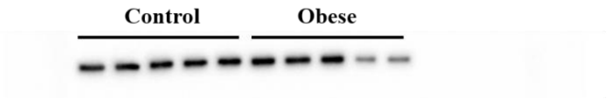


**Mitochondrial transcription factor A (Mitochondrial) (Males)**

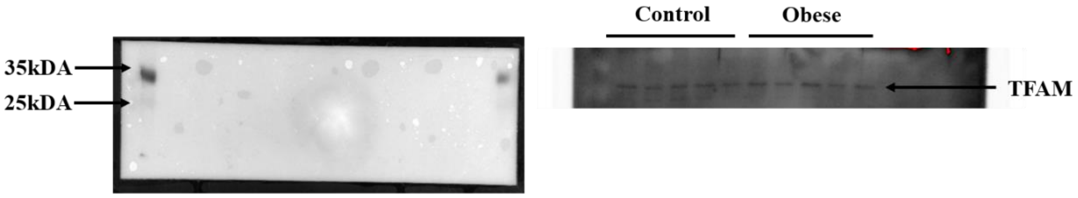


**β-tubulin (Males)**

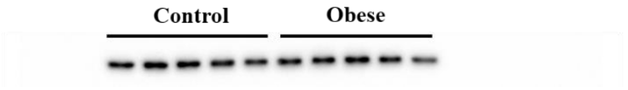

**Total OXPHOS (Complex I) (Females)**

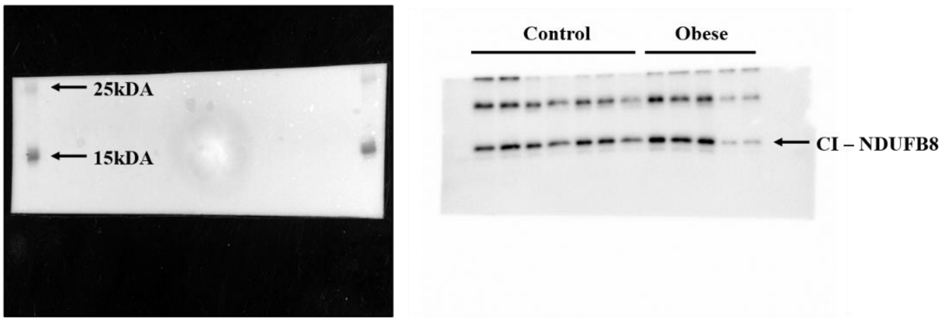


**β-tubulin (Females)**

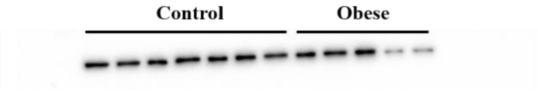


**Total OXPHOS (Complex II to V) (Females)**

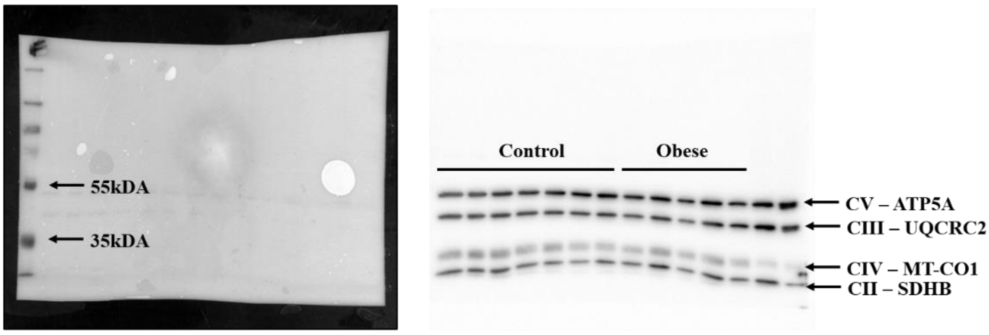


**β-tubulin (Females)**

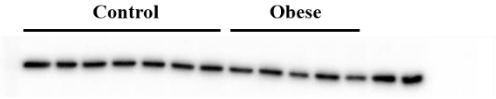

**Total OXPHOS (Complex I) (Males)**

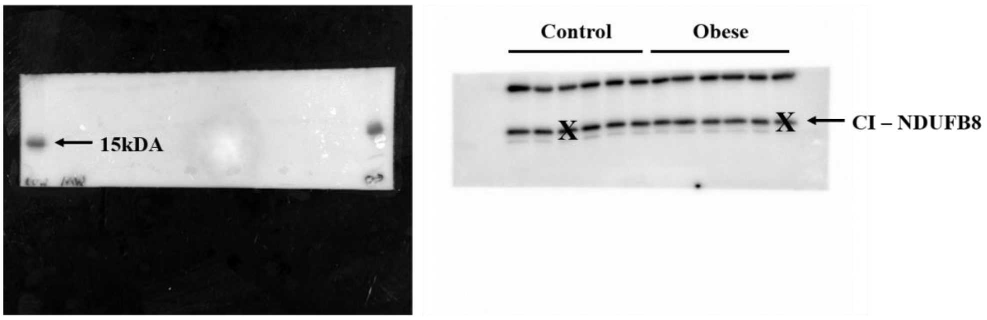


**β-tubulin (Males)**

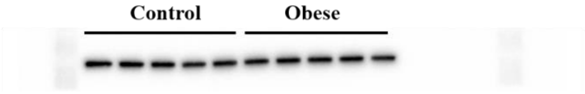


**Total OXPHOS (Complex II to V) (Males)**

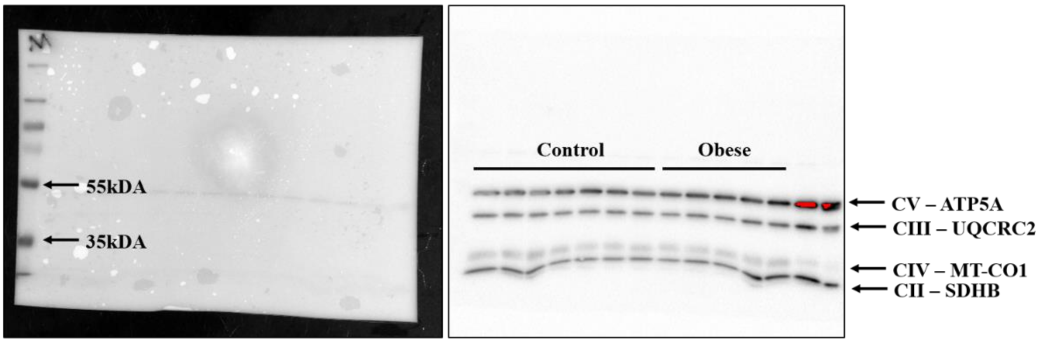


**β-tubulin (Males)**

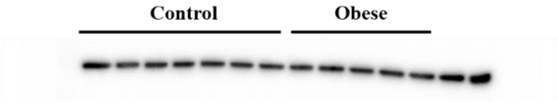

### Syntaxin 1a (Females)

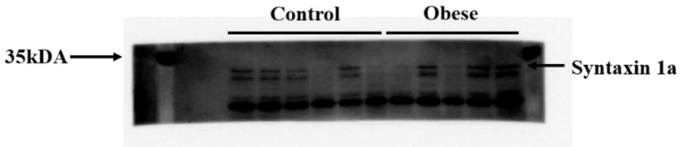


### β-tubulin (Females)

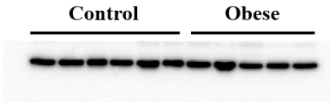


### Syntaxin 1a (Males)

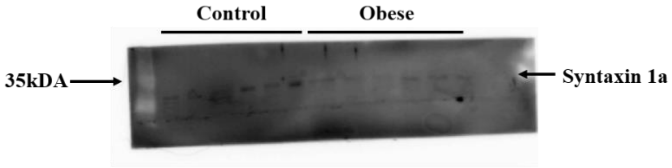


### β-tubulin (Males)

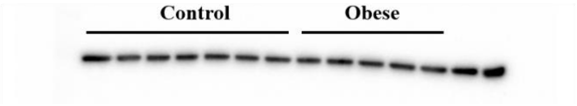


### Syntaxin binding protein 1a

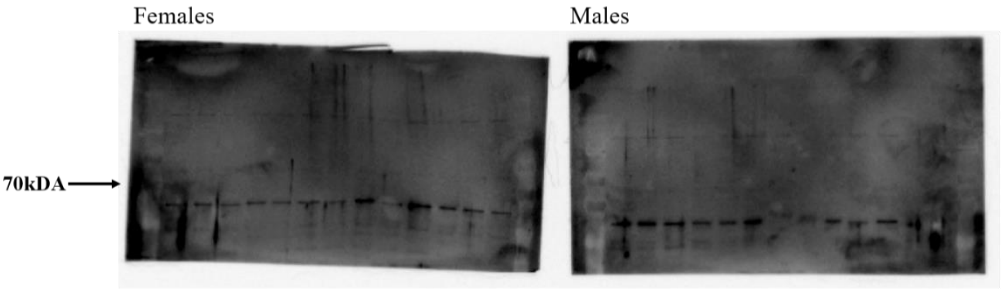


### β-tubulin

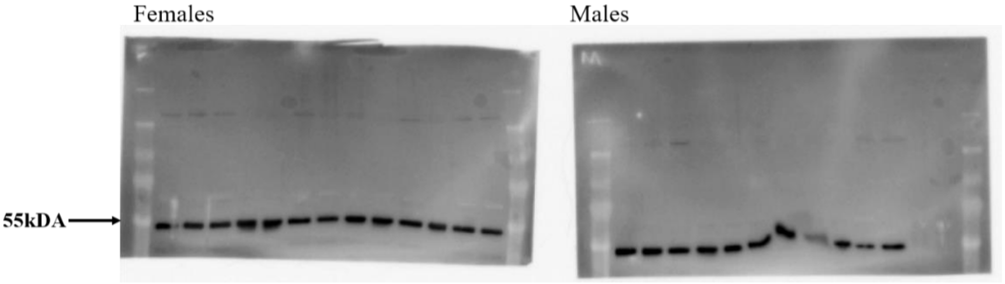

**Estrogen receptor α (Females)**

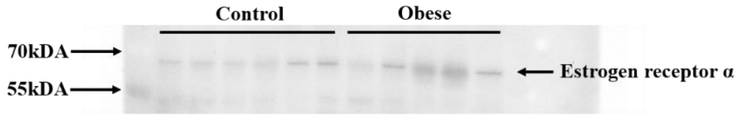


**β-tubulin (Females)**

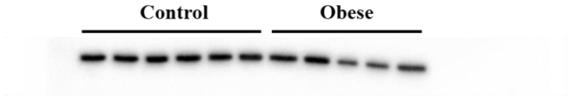


**Estrogen receptor α (Males)**

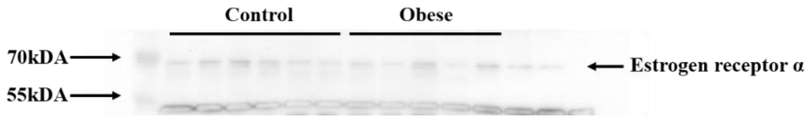


**β-tubulin (Males)**

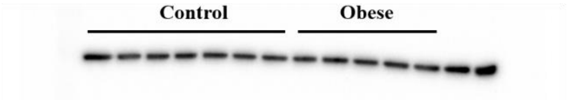


**Cleaved caspase 3 (Females)**

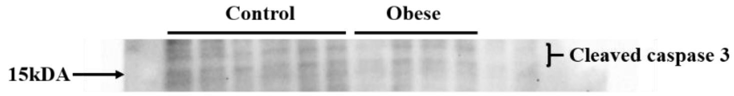


**β-tubulin (Females)**

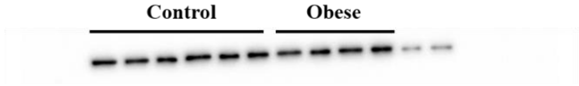


**Cleaved caspase 3 (Males)**

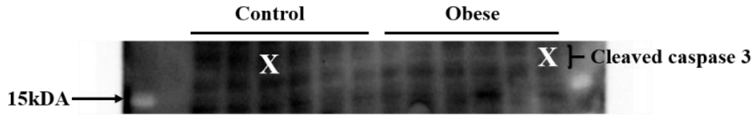


**β-tubulin (Males)**

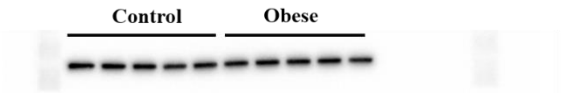

**Bax (Females)**

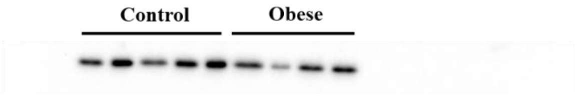


**β-tubulin (Females)**

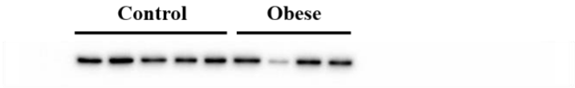


**Bax (Males)**

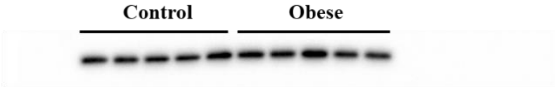


**β-tubulin (Males)**

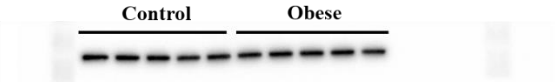


**Bcl-2 (Females)**

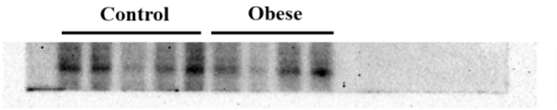


**β-tubulin (Females)**

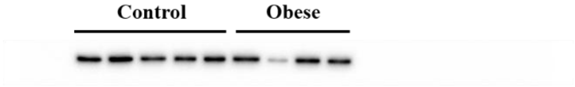


**Bcl-2 (Males)**

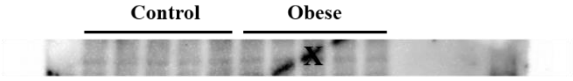


**β-tubulin (Males)**

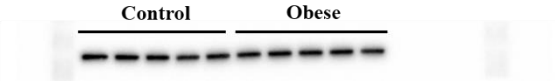
